# Protein design for evaluating vaccines against future viral variation

**DOI:** 10.1101/2023.10.08.561389

**Authors:** Noor Youssef, Sarah Gurev, Fadi Ghantous, Kelly Brock, Javier Jaimes, Nicole N. Thadani, Ann Dauphin, Amy C. Sherman, Leonid Yurkovetskiy, Daria Soto, Ralph Estanboulieh, Ben Kotzen, Pascal Notin, Aaron W. Kollasch, Alexander A. Cohen, Sandra E. Dross, Jesse H. Erasmus, Deborah H. Fuller, Pamela J. Bjorkman, Jacob E. Lemieux, Jeremy Luban, Michael S. Seaman, Debora S. Marks

## Abstract

Recurrent waves of SARS-CoV-2 infection, driven by the periodic emergence of new viral variants, highlight the need for vaccines and therapeutics that remain effective against future strains. Yet, our ability to proactively evaluate such therapeutics is limited to assessing their effectiveness against previous or circulating variants, which may differ significantly in their antibody escape from future viral evolution. To address this challenge, we develop a deep learning method to predict the effect of mutations on fitness and escape from neutralizing antibodies. We use this model to engineer 83 unique SARS-CoV-2 Spike proteins incorporating novel combinations of up to 46 amino acid changes relative to the ancestral B.1 variant. The designed constructs were infectious and evaded neutralization by nine well-characterized panels of human polyclonal anti-SARS-CoV-2 immune sera (from vaccinated, boosted, bivalent boosted, and breakthrough infection individuals). Designed constructs on contemporary SARS-CoV-2 strains displayed similar levels of antibody neutralization escape and similar antigenic profiles as variants seen subsequently (up to 12 months later) during the COVID-19 pandemic despite differences in exact mutations. Our approach provides targeted datasets of antigenically diverse escape variants for an early evaluation of the protective ability of vaccines and therapeutics to inhibit not only currently circulating but also future variants. This approach is generalizable to other viral pathogens.

## Introduction

The emergence of viral variants that evade protective immunity from prior infections, vaccines, and therapeutics is a challenge for the control of viral spread. This issue has come to the forefront during the COVID-19 pandemic. While SARS-CoV-2 vaccines and therapeutics have mitigated the severity of COVID-19, the effectiveness of many interventions has been progressively eroded by new variants of SARS-CoV-2. For instance, the FDA authorizations of all monoclonal antibody therapies have been revoked based on their loss of efficacy against emerging variants of concern (VOCs)(*1*), and the ability of first-generation vaccines and boosters to prevent infection has waned against subsequent variants(*2*, *3*). Furthermore, immunity elicited by the BA.4/5 bivalent mRNA booster vaccines is declining against the most recent variants, such as XBB, CH.1.1, and BA.2.86(*4*), leading to annually updated booster vaccines including the latest variants of concern.

Current preclinical protocols for evaluating the effectiveness of vaccines and therapeutics primarily assess neutralization potency of antisera against previous or currently circulating SARS-CoV-2 strains, or against SARS-like betacoronaviruses (sarbecoviruses) with distant phylogenetic relationships(*5–10*). These assessments, although valuable, serve as proxies for anticipating protection against future viral evolution, which is likely to diverge substantially from previous variants or related viruses. Ideally, more direct methods for anticipating immune-evasive strains can be used to generate panels of viral protein sequences against which the breadth and potency of vaccine-elicited neutralizing antibodies and therapeutics can be assessed (Fig 1A).

**Fig. 1:**
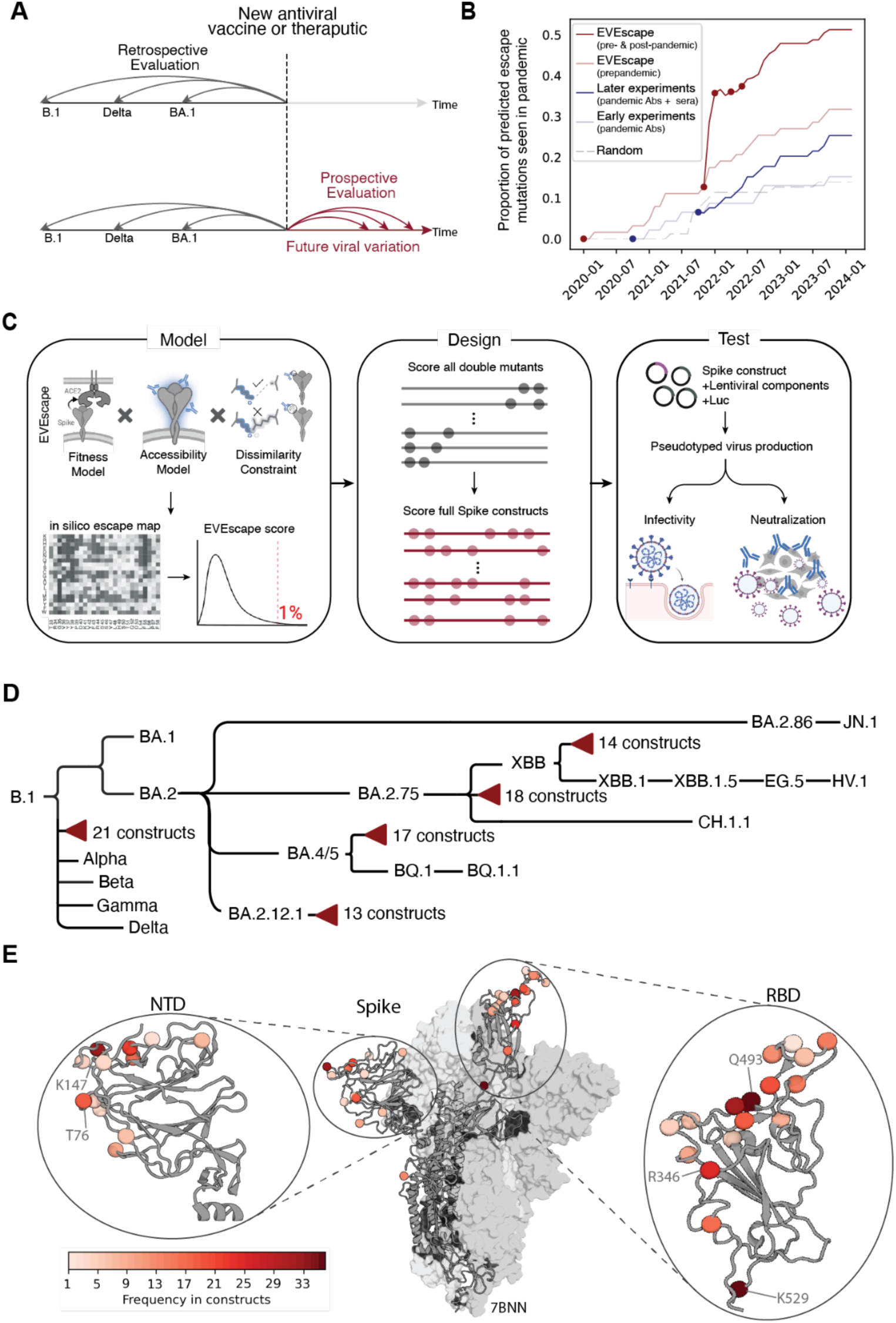
Proactive evaluation of vaccines and therapeutics using computationally designed Spike constructs. (**A**) Traditional vaccines or antiviral therapeutics evaluation are often performed retrospectively against past variants or evolutionarily related viruses. We propose a proactive evaluation approach where the immunity elicited by vaccines or therapeutics are tested against designed variants computationally predicted to foreshadow future viral variation. **(B)** Proportions of escape mutations seen more than 1000 times in GISAID as predicted by EVEscape (with and without SARS-CoV-2 sequences), mutational scan experiments (earlier and later in the pandemic), or randomly selected mutations in the RBD. Escape mutations were defined as those in the top 5% of escape scores for both the computational and experimental methods. EVEscape using pre-pandemic sequences (light red) anticipates pandemic variation significantly better than mutational scan experiments using antibodies and sera available 10 months(*11*) (light blue) or 17 months(*11*, *24*–*30*) (dark blue) into the pandemic. EVEscape performance improves as pandemic sequences are incorporated into model training. Red points indicate dates when EVEscape was updated to include sequences seen up until that date (four updates). This analysis focuses on nonsynonymous RBD mutations that are a single nucleotide distance away from the ancestral sequence. Computational predictions on full Spike are in Supp Fig2. Parts of figure created with Biorender. (**C**) Outline of computational algorithm used to design Spike proteins. All single mutations are first evaluated for their escape potential using the EVEscape model. Then, all possible double mutants combining the top 1% of highest escape single mutants are scored. Double mutants are then combined and scored, resulting in multi-mutant Spike designs. The designed constructs were then tested for their infectivity and neutralization sensitivity using pseudotyped virus assays. (**D**) Cladogram of variants of concern and designed constructs (red triangles). Branch lengths are representative of the order of variant emergence. (**E**) Designed mutations across all 83 designed constructs mapped onto a representative Spike 3D structure (PDB ID: 7BNN). Coloring indicates the frequency with which a given residue was mutated in the designed constructs.

Several experimental approaches have been developed to predict SARS-CoV-2 mutations that evade the immune response, including pseudovirus assays and high-throughput deep mutational scans(*9–12*). High-throughput approaches have been used to measure the effects of all single mutations in the receptor binding domain (RBD)(*2*, *3*, *11*, *13*) and, more recently, experiments assaying multi-mutant full Spike proteins were achieved by combining mutations frequently observed during the pandemic(*12*). However, there are challenges with using these experimental approaches for foreshadowing viral immune evasive mutations. Firstly, evolving viral strains tend to include mutations throughout the Spike protein, not only in the RBD(*14*). Second, the emerging variants often include novel mutations that evade contemporary immune responses and have not been seen before during the pandemic. *In vitro* passaging in the presence of plasma or monoclonal antibodies presents an alternative experimental approach for identifying viral escape mutations(*9*, *10*). Nevertheless, these approaches are difficult to perform at a large scale since it is impractical to exhaustively assess immune escape mutations across the diverse exposure histories and interventions available. An important drawback to most experimental approaches is that only a modest fraction of all possible multi-mutant proteins can be assayed, and these approaches cannot be exploited until after immune sera or monoclonal antibodies are available.

Deep generative models have the potential to address these challenges by rapidly assessing large numbers of multi-mutant sequences, before immune sera are available. Computational models trained on natural sequences across evolution accurately predict mutation effects across various proteins by capturing constraints, including complex dependencies among residues(*15–20*). Recently, EVEscape(*21*), a modular framework incorporating fitness effect predictions from a Baysien variational autoencoder with structural and general biophysical properties, was developed which quantifies the potential of viral mutations to evade antibodies. EVEscape, trained on sequences available prior to the onset of the COVID-19 pandemic from distantly related viruses, significantly outperformed previous computational methods and performed on par with experimental methods at anticipating RBD mutations observed during the COVID-19 pandemic(*21*). In this work, we extend the EVEscape framework to design a panel of SARS-CoV-2 Spike proteins representing future antigenic evolution at different time points throughout the pandemic (Fig 1D). We focus on the Spike protein since it is the target of most neutralizing antibodies that confer protective immunity(*22*). Our generative model does not rely on high-throughput experimental scans or on antigen-antibody structures. It is therefore generalizable to other viruses and is applicable on day-zero of a viral outbreak. We demonstrate that using EVEscape to design sets of sequences with high escape propensity presents a novel approach for evaluating the efficacy of vaccines and therapeutics against both existing and future viral variants.

## Results

A limitation in current vaccine and therapeutic evaluation approaches is that elicited protection is assessed against previous or currently circulating variants of a virus. We propose a novel approach for proactively assessing the effectiveness of vaccines and therapeutics in protecting against rapidly evolving viral variants by (1) predicting proteins that antigenically resemble future viral strains, (2) generating the proteins in biosafety level 2 conditions and (3) evaluating their escape potential from therapeutics or vaccine-elicited sera. To demonstrate this, we used variant-specific EVEscape models to design multi-mutant Spike constructs on five different VOC backgrounds: B.1 (ancestral variant with D614G), BA.4/5, BA.2.12.1, BA.2.75, and XBB.

EVEscape models the probability of an antibody escape mutation based on three constraints: the mutation’s impact on viral fitness, the accessibility of the mutated residue to antibodies, and its potential to disrupt antibody binding based on biophysical differences between the wildtype and mutant amino acid. The fitness constraint is learned using EVE(*16*), a variational autoencoder trained on distantly related protein sequences. The accessibility component is learned from three-dimensional conformations of the Spike trimer (without antibodies) and is calculated as the negative weighted residue-contact number. The dissimilarity component is computed using differences in hydrophobicity and charge between the wildtype and mutated amino acid.

EVEscape trained on data available prior to the onset of the COVID-19 pandemic (day zero) better predicts mutations observed during the pandemic than early mutational scanning experiments(*11*) (performed 10 months into the pandemic) — approximately 30% of escape mutations predicted using a pre-pandemic EVEscape model were subsequently observed >1000 times in the pandemic(*21*), compared to ∼15% of escape mutations predicted by early mutational scans(*11*) (Fig1B). Escape mutations were defined as those in the top 5% of the escape score distributions for each of the computational and experimental methods. Moreover, incorporating pandemic sequences into EVEscape training (Fig 1B, Supp Fig S1) further improves predictions – approximately 50% of predicted escape mutations from a model trained on pre-pandemic and pandemic sequences seen up to July 2022 were seen >1000 times in GISAID(*23*) to date. Note that while 7,049 SARS-CoV-2 Spike sequences were included in the latest EVEscape model, the sequences are highly similar such that the effective number of pandemic sequences was ∼15 (Supp Table S1). This enables designs on later pandemic variants to have more targeted selection of mutations that depend on their latest circulating variant.

To design multi-mutant Spike proteins, for each of the five starting background VOC spike sequences, we first scored all possible single amino acid mutations (∼121,000 for each VOC; Fig 1C) and identified mutations with high escape potential (top 1% EVEscape scores). Then, we calculated the escape potential of all high scoring double mutants (∼140,000 for each VOC; Fig 1C). Finally, we combined the double mutants to create full Spike constructs with higher-order mutations (Fig 1D, Supp Fig S2). For the B.1 variant we developed two models. The first was trained only on sequences available prior to the onset of the COVID-19 pandemic, including Spike sequences from other coronaviruses (total of 4,577 non-SARS-CoV-2 Spike sequences; Supp Table S1). The second model was updated to include SARS-CoV-2 Spike sequences available in May 2022 (1,291 SARS-CoV-2 sequences which were highly similar and had an effective number of ∼9 sequences; Supp Table S1). For all other variants, we updated the EVEscape model to include pandemic sequences that occurred up until the time of emergence of that particular variant(*21*) (Supp Table S1-S2).

Using this approach, we designed 83 unique Spike proteins predicted to escape polyclonal sera across mutational depths, including both rare and common mutations seen in GISAID (Supp Table S3, Supp Fig S3). Constructs contained up to 10 novel combinations of mutations relative to the nearest VOC, and up to 46 mutations relative to the ancestral B.1 strain (Supp Table S4). In total, 37 unique mutations, across 30 positions, were designed in different combinations and on different Spike backgrounds. Consistent with previous EVEscape predictions for escape mutations, most designed mutations were located in primary antigenic regions observed in the pandemic(*21*, *22*) — 17 (57%) of the mutated residues were in the RBD and 12 (40%) were in the NTD (Fig 1E).

### Computationally-designed multi-mutant Spikes maintain infectivity in vitro

A challenge with generating multi-mutant proteins for proactive vaccine and therapeutic evaluation is that most mutated proteins lose functionality as they deviate further away from the wildtype protein. For instance, randomly mutating the SARS-CoV-2 receptor binding domain (RBD) using error-prone PCR resulted in only ∼2% of sequences with eight or more mutations being expressed(*13*) (Fig 2A). Note that this is likely an overestimate of the proportion of viable full Spike sequences since mutations that are stable and well-folded in the isolated RBD may not be accommodated in the Spike trimer. Similarly, the proportion of functional Spike proteins achieved by combining mutations observed during the pandemic(*12*) decreased progressively with further mutations from the wildtype(*12*) (Fig 2A). Together, these results highlight the importance of context-dependence of mutations where specific mutations may be viable in one background but not in another.

**Fig. 2:**
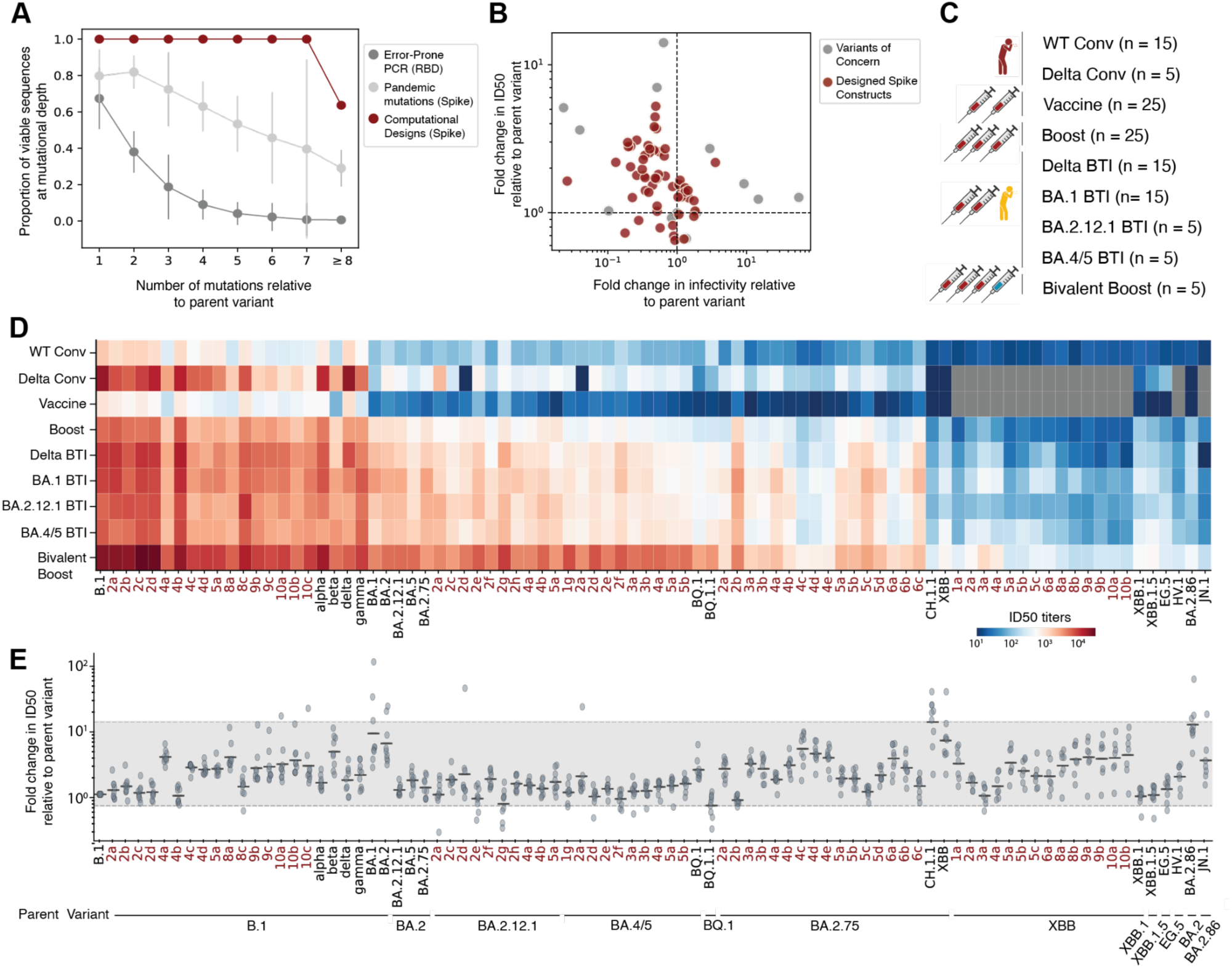
Designed constructs have similar levels of neutralization escape as variants of concern. (**A**) Proportion of viable sequences across mutational depths. Designed constructs are more likely to be viable than sequences resulting from error-prone PCR within the RBD (assayed for expression(*13*)) or from combinations of frequently observed pandemic mutations in Spike (assayed for infectivity(*12*)). For the experimental datasets, the proportion of functional sequences was estimated by performing 10,000 samples of n sequences, where n is the number of computationally designed sequences, at a given mutational depth. Error bars represent one standard deviation. (**B**) Relationship between infectivity and neutralization. Most SARS-CoV-2 variants and designed constructs that demonstrated higher levels of escape were less infectious relative to their parent variant. (**C**) Summary of serum panels from individuals who were either convalescent (Conv.), vaccinated, boosted, or experienced breakthrough infection (BTI). A total of 23 serum pools were created by grouping 5 individual patient samples per pool. (**D**) Neutralizing ID50 titers for each VOC and designed construct across nine sera panels. Reported are the geometric mean titers across all sera pools within a panel. Gray boxes indicate variants that were not tested in that serum panel. Designs are highlighted in red text. (**E**) Fold change in ID50 relative to parent variant for each VOC and background variant for each design. Each point represents the geometric mean fold change for a given sera panel (nine panels in total). Lines indicate geometric mean across sera panels. The gray band indicates the observed range of changes in neutralization sensitivity across variants of concern. Designs are highlighted in red text.

To test infectivity of the designed Spike sequences, we used lentiviral-based pseudoviruses in single-round biosafety-level-2 infection assays, as previously described(*31*). Among the 83 designed constructs, 90% (75/83) were infectious (Fig 2A, Supp Table S5, Supp Fig S4A). The proportion of functional Spike proteins achieved by computational design was consistently higher than that achieved by combining mutations observed during the pandemic(*12*). Our model provides consistent performance across mutational depths, highlighting the advantage of using deep generative models for designing multi-mutant Spike constructs and facilitating early vaccine and therapeutic evaluation.

Four of the eight designed constructs that were not infectious included the L452R, F490R, Q493S, and Q498Y mutations. Note that other constructs containing some of these mutations individually, or in pairs, retained functionality and were infectious. For example, constructs designed on BA.2.12.1 containing the mutation pairs L452R and Q498Y (construct 2a), L452R and Q493S (2b), and F490R and Q498Y (7a) were infectious (Supp Fig S5A). We hypothesize that the proximity of L452, F490, and Q493 in the three-dimensional structure leads to a steric clash and might contribute to the lack of infectivity. This is consistent with the observation that no variants have been detected during the pandemic that contain such closely-spaced triplet mutations (Supp Fig S5B-C). The remaining four non-infectious constructs were designed on the B.1 background using the pre-2020 model. They all included the R403T mutation which was shown to disrupt ACE2 binding(*32*), likely explaining their lack of infectivity. These findings provide valuable insights into the enhancement of the design algorithm.

To further evaluate the impact of designed mutations on functional activity, we compared the infectivity of each variant relative to its parent variant (Fig 2B). The initial Omicron variants BA.1 and BA.2 had over 30 Spike mutations relative to B.1, concurrent with a 43.5-fold and a 25.4-fold decrease in infectivity relative to B.1 (Fig 2B, Supp Fig S4, P < 0.05, Mann-Whitney rank-sum test). Subsequent BA.2 sub-lineages, BA.2.12.1 (2 additional Spike mutations) and BA.2.75 (6 additional Spike mutations) recovered infectivity with a 5.6-fold and a 28.9-fold increase relative to parent variant BA.2 (Supp Fig S4, Fig 2B, P < 0.05, Mann-Whitney rank-sum test). Designed constructs retained comparable levels of infectivity relative to the background variant on which they were designed despite containing up to 10 mutations. On average, designed constructs had a 1.2-fold reduction in infectivity relative to their parent variant (Supp Fig S4). Collectively, these results underscore the efficacy of this approach in producing multi-mutant Spike constructs that retain *in vitro* infectivity.

### Spikes designed on early variants exhibit neutralization resistance similar to subsequent SARS-CoV-2 variants

The continued efficacy of vaccines and therapeutics in part depends on the ability of elicited antibodies to retain the capacity to neutralize newly emerging viral variants. To assess the relevance of the designed Spike constructs as representative of future viral variation, we performed pseudovirus neutralization assays to evaluate sensitivity profiles when tested against nine panels of human polyclonal serum pools representative of diverse SARS-CoV-2 immune histories: (1) individuals with convalescent B.1 infection prior to the availability of vaccines, (2) unvaccinated individuals with convalescent Delta infection, (3) recipients of two primary doses of mRNA-1273 or BNT162b2 vaccines, (4) individuals who received three doses of an mRNA vaccine, (5) vaccinated individuals administered a BA.4/5 bivalent booster (mRNA-1273 or BNT162b2), and vaccinated patients who experienced breakthrough infections (BTI) with (6) Delta, (7) BA.1, (8) BA.2.12.1, or (9) BA.4/5 variants (Fig 2C, Supp Table S6). Serum panels were tested against pseudoviruses expressing 66 of the designed Spike constructs, 20 SARS-CoV-2 variant Spike proteins (B.1, Alpha, Beta, Delta, Gamma, BA.1, BA.2, BA.2.12.1, BA.2.75, BA.4/5, BQ.1, BQ.1.1, XBB, XBB.1, XBB.1.5, CH.1.1, EG.5, HV.1, BA.2.86, JN.1), and the SARS-CoV-1 Spike protein (Fig 2D, Supp Table S7).

To characterize the degree of immune-evasion observed throughout the pandemic, we compared the half-maximal neutralizing antibody titers (ID_50_) for each variant relative to the parent variant from which it evolved (Fig 2E). As expected, most variants were less susceptible to neutralization (had higher antibody escape) compared to the parental variant that preceded it, with the exception of BQ.1.1, XBB.1 and XBB.1.5 variants which had a comparable geometric mean ID_50_ titer across sera panels to their parent variants BQ.1, XBB, and XBB.1, respectively (P > 0.2, Wilcoxon rank-sum test). The XBB (7.2-fold, P < 0.01, Wilcoxon rank-sum test) and CH.1.1 (14.2-fold, P < 0.01, Wilcoxon rank-sum test) variants displayed the highest level of escape relative to their parent variant BA.2.75. On average emerging variants exhibited a 3.9-fold reduction in geometric mean ID_50_ titers relative to their parent variant, and a range from 0.67 to 14.2 (gray region, Fig 2E). Generally, variants that had higher levels of antibody escape tended to be less infectious than their parent variant (Fig 2B).

Pseudoviruses expressing designed Spikes had comparable levels of antibody escape relative to their parent variants as has been observed for SARS-CoV-2 variants that evolved throughout the pandemic. On average, designed Spikes had a 1.9-fold reduction (range of 0.5 to 5.31) in geometric mean ID_50_ titer compared to the parent variant on which they were designed (Fig 2E). More specifically, designs on specific backgrounds had similar levels of neutralization escape as SARS-CoV-2 variants evolved from those same backgrounds. For instance, the B.1-4a construct designed construct, containing the K147N, S494R, F490R, and R683N mutations on the B.1 background, had a 3.9-fold reduction in geometric ID_50_ titers, which is higher than the level of escape observed for the alpha, delta, and gamma variants (1.5, 1.7, and 2.0-fold reduction relative to B.1 respectively) and similar to the escape seen in the beta variant (4.8-fold reduction relative to B.1). Similarly, the BA.4/5-2a designed construct, containing the R346T and S494R mutations on the BA.4/5 background, had a 1.9-fold reduction in geometric ID_50_ titers, which is comparable to the level of escape observed for BQ.1 (2.4-fold reduction relative to BA.4/5). The BA.2.75-4c design, containing G339D, L452R, Q493R, and K529L, had a 5.31-fold reduction in geometric mean ID_50_ titer relative to BA.2.75, slightly lower than the level of escape observed by XBB (7.2-fold reduction in geometric mean titers relative to BA.2.75). Taken together these results demonstrate that Spike constructs designed using EVEScape exhibit a similar level of antibody escape as variants that have naturally evolved under immune pressure throughout the pandemic and can therefore serve as useful proxies for future SARS-CoV-2 evolution and guide early evaluation of vaccines and therapeutics.

### Designed variants foreshadow future antigenic evolution

Thus far we have demonstrated that our designed Spike constructs are comparable to SARS-CoV-2 variants in both infectivity and neutralizing antibody escape. We then sought to determine whether these constructs accurately reflect the antigenic evolution observed throughout the pandemic. To investigate this, we constructed antigenic maps from the neutralization data using a multidimensional scaling (MDS) algorithm to characterize the space of neutralizing antibody responses across variants (Fig 3A). The resulting antigenic cartography provides a consistent two-dimensional representation such that distance between any pair of variants is related to the similarity in which they are neutralized by the different sera pools that represent populations with varied infection or vaccination histories (Supp Fig S6).

**Fig. 3:**
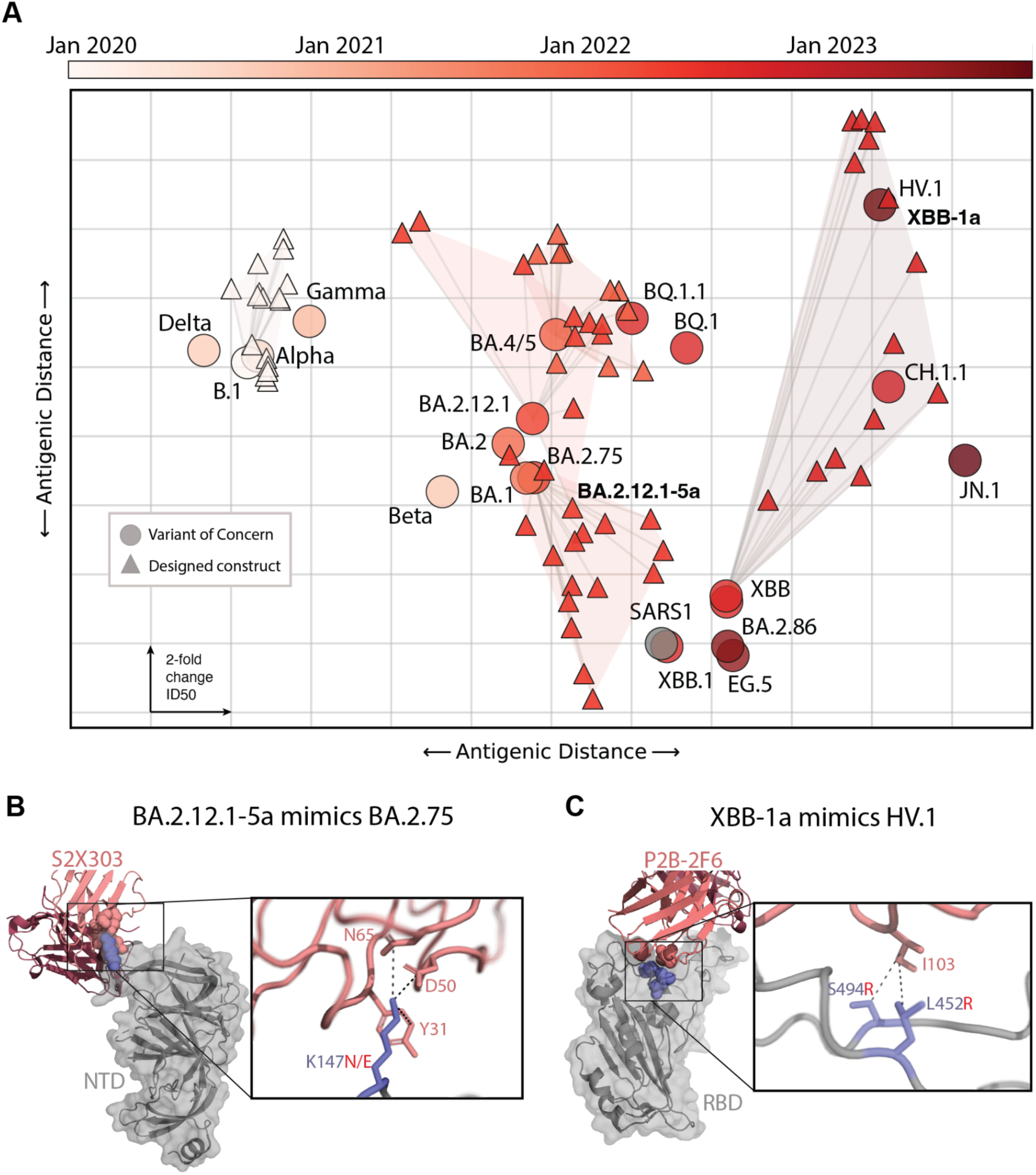
Designed constructs foreshadow future SARS-CoV-2 evolution. (**A**) Antigenic cartography depicts a two-dimensional representation of the pairwise antigenic distance between each variant of concern (circles) or designed construct (triangles), as dictated by how each is neutralized across the 23 serum panels. The dimensions of each grid box represents a 2-fold change in neutralizing ID50 titers. Variants are colored based on the month of emergence since the onset of the pandemic. Constructs designed on earlier variants (triangles of lighter colors) foreshadow the antigenicity of later pandemic variants (circles of darker colors), as indicated by proximity in the antigenic map. (**B**) Design BA.2.12.1-5a mimics the neutralizability of later pandemic variant BA.2.75 as seen in Fig3A. Each contains a mutation at K147 (N in the design, E in the later variant). The impact of K147 mutations can be seen for NTD antibody S2X303, whose epitope partially overlaps the NTD antigenic supersite(*33*), with interactions to N65, D50, and Y31 on the antibody (PDB ID: 7SOF). (**C**) A cluster of many designs containing L452R (XBB-8a,9a,9b,10a,10b) closely match the neutralizability of the later pandemic variant HV.1 (also containing L452R) as seen in Fig3A. The XBB-1a (XBB + S494R) design almost exactly mimics the neutralizability of the later HV.1 (XBB + Q52H + F157L + G252V + L452R + F456L) variant. The impact of the L452R and S494R mutations can be seen in Wuhan convalescent RBD class II antibody P2B-2F6(*34*), where mutation to an arginine of either S494 or L452 (which both contact I103 on the antibody) result in escape(*2*) (PDB ID: 8DCC).

Constructs designed on earlier variants consistently exhibited an antigenic resemblance to variants that emerged later in the pandemic. For example, constructs designed on BA.2.75 were antigenically similar to the XBB lineages which emerged nine months later (Fig 3A, shaded area). Similarly, constructs designed on the XBB variant exhibited antigenic responses akin to the CH.1.1 and HV.1 variants, which emerged three and 12 months later (Fig 3A, shaded area). This finding holds for all other rounds, with designs on BA.2.12.1 foreshadowing BA.4/5, designs on BA.4./5 foreshadowing BQ.1.1, and designs on WuG foreshadowing the Alpha and Gamma variants (Fig 3A).

Critically, designs are able to mimic future escape from antibodies elicited by particular vaccination or infection histories even while testing a relatively limited number of constructs. For instance, design BA.2.12.1-5a mimics the neutralizability of later pandemic variant BA.2.75 (Fig 3A), while each contains a different mutation to K147 (N in the design, E in the later variant). The impact of K147 mutations can be seen for NTD antibody S2X303, whose epitope partially overlaps the NTD antigenic supersite(*33*), with many amino acids likely causing escape due to its highly constrained three antibody contacts (Fig 3B). Secondly, a cluster of many multi-mutant designs containing L452R (XBB-8a,9a,9b,10a,10b) closely match the neutralizability of the later pandemic variant HV.1 (also containing L452R). Yet even more strikingly, the XBB-1a design with only a single mutation to XBB (S494R, which neighbors L452R) almost exactly mimics the neutralizability of the later HV.1 variant. The similar impact of these neighboring mutations can be seen in Wuhan convalescent RBD class II antibody P2B-2F6(*34*), where mutation to an arginine of either S494 or L452 (which both contact I103 on the antibody) result in escape(*2*) (Fig 3C). This example also showcases how many designs exhibited outsized effects on antigenicity relative to their number of mutations (Supp Fig S6), especially mutations to the more neutralizing subregions of Spike, the receptor binding motif and the NTD antigenic supersite(*21*, *35*) (the location of many designed mutations, despite this not being information the model is trained on). Moreover, XBB-1a (XBB + S494R) and HV.1 (XBB + Q52H + F157L + G252V + L452R + F456L), have notable escape from BA.1 BTI sera compared to other XBB designs and more so than differences for other sera panels (Supp Table S7). Later Omicron strains are thought to have evolved L452R specifically to maximize immune evasion after infection with BA.1(*3*). Collectively these findings underscore the model’s capability to design constructs with antigenic profiles akin to those of future variants and showcase its potential in predicting antigenic evolution using only data available at the time of VOC emergence.

### Evaluating mRNA vaccine elicited neutralizing antibodies

To demonstrate the utility of the proposed approach for proactive evaluation of vaccine candidates, we showcase how designed variants could have been utilized to evaluate the ability of neutralizing antibodies elicited by the BA.4/5 bivalent mRNA booster vaccines to protect against future evolving variants. Considering variants circulating in the four months prior to the implementation of the bivalent booster (BA.2.75, BQ.1, BQ.1.1, and XBB), the geometric mean ID_50_ titers for serum panels representative of bivalent boosting (4 shots) were 1,931 (Fig 4A, Supp Table S7). Constructs designed on the variants circulating at the time of the booster vaccination campaign (designs on BA.2.75 and XBB) demonstrated a range of antibody escape with ID_50_ ranging from 193 to 4,029, and had an average 2.6-fold decrease in geometric mean titers relative to the VOCs present prior to the bivalent vaccine (P = 0.025, Wilcoxon rank-sum test, Fig 4A). Notably, levels of antibody escape seen in variants that evolved in the four months after the implementation of the bivalent boosters XBB.1, XBB.1.5 and CH.1.1 (ID_50_ titers of 538, 587, and 308 respectively) fall within the distribution of titers observed in the designed constructs (Fig 4A). Therefore, designed constructs on variants circulating at the time of the bivalent booster vaccination campaign could have highlighted the potential for antibody escape observed in subsequent lineages. Furthermore, eight of the designed constructs on XBB had lower ID_50_ titers than CH.1.1, demonstrating the potential for further escape from neutralizing antibodies elicited by bivalent booster vaccines.

**Fig. 4:**
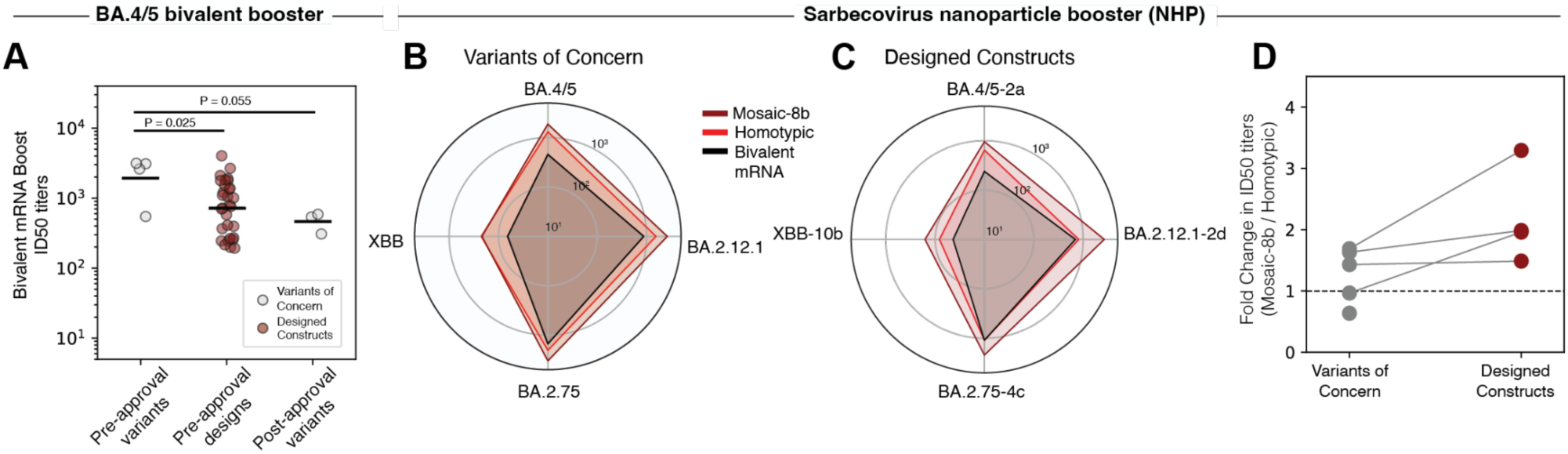
Evaluating the breadth and potency of neutralizing antibodies elicited by vaccines using designed Spike constructs. (**A**) Neutralizing ID50 titers from individuals who received three doses of mRNA-1273 or BNT162b2 and a fourth dose of the BA.4/5 bivalent vaccine. Lines indicate geometric mean across variants or designs. The BA.4/5 bivalent booster vaccine was evaluated on its ability to neutralize variants circulating in the four months prior to the creation of the booster vaccines (BA.2.75, BQ.1, BQ.1.1, and XBB), which was approved on Aug 31, 2022. Designed Spike constructs on the background of variants circulating prior to the vaccine foreshadowed the levels of antibody escape observed against variants that emerged following the bivalent booster (XBB.1, XBB.1.5, CH.1.1). Significance assessed using the Wilcoxon rank-sum test. (**B**) Neutralizing potency of sera collected from non-human primates primed with a mixture of nucleic acid vaccines and boosted with either a bivalent mRNA (n=4), a homotypic nanoparticle (n=5), or a mosaic 8b nanoparticle (n=5). Radar charts represent for each variant the logarithm of the geometric median ID50 titers. (**C**) Radar charts represent the logarithm of the geometric median ID50 titers for each designed construct. Designed constructs can aid in selection of a nanoparticle booster as they more easily show a difference between the mosaic-8b and homotypic nanoparticles. (**D**) Relative neutralization sensitivity of a given variant to the mosaic-8b boost sera relative to the homotypic boost sera. Higher values indicate the mosaic-8b vaccine better neutralizes a given variant. Lines connect designed constructs to the variant they were designed on.

### Evaluating nanoparticle vaccine elicited neutralizing antibodies

Next, we examined the utility of our approach for evaluating neutralizing antibody responses elicited by candidate nanoparticle-based vaccines. Cohen *et al*(*7, 8*) recently developed two RBD nanoparticle vaccines: Mosaic-8b nanoparticles displaying RBDs from eight different sarbecoviruses, and a homotypic nanoparticle displaying the RBD from the Beta SARS-CoV-2 variant. Both nanoparticle vaccines elicited neutralizing antibodies against SARS-CoV-2 variants when tested in mouse and non-human primate (NHP) animal models(*7*, *8*). To evaluate the neutralizing potential of these booster vaccines against future SARS-CoV-2 evolution, we examined differences in neutralization activity across five SARS-CoV-2 variants (WuG, BA.2.12.1, BA.2.75, BA.4/5, and XBB) as well as four designed Spike constructs BA.5-2a, BA.2.12.1-2d, BA.2.75-4c, and XBB-10b; Supp Table S8). We chose the design on each background that had the highest level of antibody escape across patient sera.

Fourteen NHPs were primed with a mixture of nucleotide vaccines and boosted with either a bivalent mRNA vaccine (n = 4), a mosaic-8b nanoparticle (n = 5), or a homotypic nanoparticle (n = 5) vaccine(*36*). Sera from bivalent mRNA-boosted NHPs demonstrated the lowest neutralization titers against the tested SARS-CoV-2 variants (678) compared to sera from NHPs which were boosted with the homotypic nanoparticle (1,843, P = 0.166 Wilcoxon rank-sum test) or the mosaic-8b nanoparticle (2,199, P = 0.060, Wilcoxon rank-sum test, Fig4B). Neutralizing titers were consistently lower against the designed constructs compared to the SARS-CoV-2 variants, with a 2.3-, 4.0-, and a 2.3-fold decrease in the bivalent mRNA, homotypic, and mosaic-8b sera, respectively (Fig 4C). While the level of neutralization among the SARS-CoV-2 variants was comparable between the mosaic-8b and homotypic boosted NHPs, the designed constructs showcase the improved performance of the mosaic-8b nanoparticle booster vaccine (Fig 4D). Neutralizing titers against the designed constructs BA.2.12.1-2d, BA.2.75-4c, BA.5-2a, and XBB-10b were 3.3-, 2.0-, 1.5-, and 2.0-fold higher in sera from mosaic-8b boosted NHPs compared to homotypic boosted animals. Overall, the results suggest that, as compared to bivalent mRNA booster vaccines, nanoparticle booster vaccines lead to higher neutralizing titers against potential future SARS-CoV-2 variants, and that the mosaic-8b elicits more cross-reactive antibodies compared to the homotypic nanoparticle.

## Discussion

Driven by the need to develop vaccines and therapeutics that remain effective in the face of viral evolution, nearly 400 SARS-CoV-2 vaccine candidates are currently undergoing preclinical or clinical evaluations(*37*). However, a significant challenge lies in our limited ability to predict whether these candidate vaccines will effectively combat future viral variants. We demonstrate the potential of deep learning models in predicting the escape patterns from neutralizing antibodies, enabling the generation of Spike protein constructs that anticipate the antigenic landscape of future SARS-CoV-2 variants. Panels of designed Spike constructs can be used to assess the ability of candidate vaccines and therapeutics to neutralize future viral variants.

We highlight the utility of our approach through (1) a retrospective analysis of the bivalent booster vaccine, and (2) a proactive evaluation of pan-sarbecovirus nanoparticle-based vaccines. We demonstrate that the immune-escape observed by currently circulating SARS-CoV-2 variants, such as XBB and CH.1.1, could have been anticipated at the time that bivalent booster vaccines were implemented, and that designed constructs demonstrate the potential for further immune escape. Alternatively, nanoparticle-based vaccines offer a promising vaccine modality that has the potential to elicit neutralizing antibodies with greater breadth against SARS-CoV-2 variants. The conceptual premise posits that vaccines designed to counteract sarbecoviruses with broad taxonomic distances, would elicit the production of antibodies targeting conserved regions, thus decreasing the likelihood of escape under immune pressures. We show that the homotypic and mosaic nanoparticles elicit higher neutralizing titers relative to bivalent mRNA boosters in nonhuman primates. Furthermore, antibodies elicited by mosaic nanoparticle boost vaccination resulted in higher neutralizing titers against designed constructs on various circulating SARS-CoV-2 variants, highlighting their potential advantage over homotypic nanoparticles as booster vaccines.

The approach presented here is generally applicable across pan-coronavirus vaccines and therapeutics and is applicable to other rapidly evolving viruses. An advantage of the modeling framework is its adaptability across viral families, particularly viruses with limited data, such as Lassa virus(*21*). Additionally, the design algorithm can be used to predict mutations within a particular epitope for the evaluation of individual antibody therapeutics. While antibody neutralization is likely a critical driver of initial clinical efficacy(*38*), ideally a similar approach could be developed for predicting and evaluating T-cell responses.

To date, cross-neutralization potential of pan-variant therapeutics(*5*, *39*), as well as SARS-CoV-2 boosters(*40*) and other antivirals(*41*), have been evaluated against previous or contemporary SARS-CoV-2 variants or against distantly-related sarbecoviruses(*7*, *8*). Such approaches are limited to retrospectively testing the diversity seen within a pandemic or throughout evolution, rather than proactively measuring protection against future viral strains. The vaccine and therapeutic evaluation approach presented here addresses these issues by first anticipating mutations that will escape current antibody repertoires, then generating pseudotyped viruses containing complex mutations, and ultimately evaluating the ability of vaccines and therapeutics to elicit antibodies that not only neutralize previous and circulating variants but also future viral evolution.

## Methods

### Computational Spike protein design

#### Multiple Sequence Alignments for fitness models

We designed Spike constructs on the backgrounds of five VOCs: the canonical wildtype SARS-CoV-2 Spike protein (Uniprot ID: P0DTC2) with D614G mutation, and the BA.4/5, BA.2.12.1, BA.2.75, and XBB Spike sequences. We define the pandemic lineages based on the list of mutations described in outbreak.info(*42*). For each round of construct design, we built multiple sequence alignments for Spike with both pre-pandemic sequences and pandemic Spike sequences seen before the emergence of the variant used as the background for the design. To generate the pre-pandemic alignment, we used 5 iterations of the jackhmmer iterative HMM-based homology search alignment tool(*43*), searching against the Uniref100 dataset (*44*) downloaded on March 16, 2022. To account for the growing number of pandemic sequences, we also included unique pandemic sequences seen more than 100 times in the Global Initiative on Sharing All Influenza Data (GISAID) EpiCoV database (www.gisaid.org)(23) at the time of emergence of the variant (Supp Table S1). The resulting pandemic sequences and evolutionary alignments were combined and aligned with super5(*45*), version 5.1. We remove sequences that are aligned to less than 50% of the query sequence and remove residue positions that contain more than 70% gap characters.

#### Modeling approach

We use the EVEscape model(*21*), which combines fitness, accessibility, and dissimilarity, to predict a mutant’s ability to escape antibody recognition. For the fitness (*f*) component we trained an EVE model(*16*), a Bayesian variational autoencoder (VAE), to predict the fitness scores. We down weighted redundant sequence clusters by assigning each protein sequence *i* a weight *w* (*i*) = 1/*T*, where to *T* is the number of sequences in the alignment within a given hamming distance cutoff of *t*. We combine the pre-pandemic and pandemic alignments and use a *t* value of 0.01. We estimate the relative fitness of each sequence as the log likelihood ratio between a mutated sequence and the wild type sequence. This ratio is itself approximated as the difference in Evidence Lower Bound (ELBO). We estimate ELBO values using twenty thousand Monte Carlo samples of the latent space.

For the dissimilarity (*d*) component, we use both charge and hydrophobicity(*46*) differences between each pair of amino acids and then assign each pair a dissimilarity value equal to the sum of the standard-scaled differences. For the accessibility (*a*) component, we selected Spike structures from the RCSB Protein Data Bank (PDB) representative of both “open” and “closed” configurations (PDB IDs: 6VXX, 6VYB, 7CAB, and 7BNN). Using these structures, we calculated the weighted contact number (WCN) for each residue position *i* as

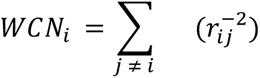

where *r_ij_* is the distance between the geometric centers of the side chain of the residues occupying sites *i* and *j*. Each position is then assigned the minimum weighted contact number across all structures. We use the negative weighted contact number as a measure of residue accessibility to antibodies.

We combine the mutation-level fitness, accessibility, and dissimilarity using a temperature scaled logistic function to get a single escape score for each individual mutation. The score of a mutation *m* is calculated as

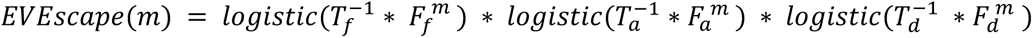

where *T_i_* is the temperature scaling and *F_i_* is the standardized vector for factor *i*. We then take the log transform of the product. We modified the temperature parameters compared to the values used previously(*21*) by increasing the contribution of the fitness component, *T_f_* = 1, *T_a_* = 8, and *T_d_* = 16.

#### Design algorithm

We designed 83 constructs on the backgrounds of five different Spike VOC sequences: B.1 (ancestral strain with D614G), BA.4/5, BA.2.12.1, BA.2.75, and XBB. For each background variant we first scored all possible single amino acid substitutions to Spike. Note that we consider amino acid mutations involving any number of nucleotide changes rather than focusing on only single nucleotide changes which are by far the most common (more than 99%) mutations seen in GISAID(*23*). To design constructs, we considered single amino acid mutations in the top 1% from our complete EVEscape score. We then generated and scored all possible double mutants based on their EVEscape score alone. Lastly, we combined highly fit double mutants and scored them to create higher order constructs, with a mix of mutations that had and had not been seen yet in the pandemic (Supp Table S4).

### Testing Infectivity

#### DNA sequence design and plasmid

We designed a pDMJ2 plasmid where Spike expression is under the control of a cytomegalovirus immediate early promoter (CMV-IE) fused to the human T lymphotropic virus type 1 (HTLV-1) 5’ UTR (*47*). Transcription termination is directed by a bovine growth hormone polyadenylation signal (bGH-PolyA). The pDMJ2 plasmid carries an origin of replication (ori) from ColE1 and an Ampicillin resistance gene. Spike sequences were inserted between the *XbaI* and *BamHI* restriction sites (Supp Table S4; Supp Fig S7).

An ancestral B.1 (ancestral variant with D614G) Spike sequence was codon optimized by Twist Biosciences with a deletion of the terminal 18 amino acids to remove the endoplasmic reticulum retention signal (*48*). This sequence was used as the base sequence on which other constructs were designed to minimize effects of nucleotide sequence on protein synthesis efficiency. Specifically, to generate Spike sequences of VOCs and designed constructs, we mutated individual codon positions to the most commonly used codon in human cells encoding the mutant amino acid. We repeated this for all mutations in a given sequence. If a restriction site was introduced, the second most common codon was used and so on.

#### Infectivity assay

HIV-1-derived virions bearing a luciferase reporter gene, and pseudotyped with SARS-CoV-2 Spike, were produced by transfection of HEK293T cells, as previously reported (*31*). The supernatant from the transfected cells was collected 72 hrs post-transfection and filtered (0.45 µM, Avantor^TM^) to remove cellular debris. Virion yield in each transfection supernatant was normalized using our in-house RT assay (*31*, *49*). Vector supernatants were then placed on HEK293T cells that were previously transduced with puromycin- and blasticidin-resistant lentivectors to stably express the human angiotensin converting enzyme 2 (ACE2) and the transmembrane serine protease 2 (TMPRSS2) (*31*), respectively. Luciferase activity on target cells was measured as an indication of transduction efficiency, using the Steady-Glo^®^ Luciferase Assay System (Promega Corporation) read on a Promega GloMax Discover machine. All plasmids were deposited to Addgene (Supp Table S4). Experiments were performed in triplicate.

#### Comparison to experimental assays

We used data from recent deep mutation scans to quantify the likelihood of obtaining functional sequences with multiple mutations compared to a wild type sequence. We analyzed the RBD data from Starr *et al*(*13*) and the full Spike data from Dadonaite *et al*(*12*). In the RBD dataset, mutations were introduced using error-prone PCR resulting in 135,386 unique mutant RBDs with up to 11 amino acid mutations. The expression of each sequence was measured as the change in mean fluorescence intensity, Δlog(MFI), relative to the unmutated SARS-CoV-2 RBD. We used sequences containing synonymous mutations (labeled viable) and sequences with premature stop codons (labeled nonviable) to train a logistic regression for classifying missense variants as either viable or nonviable. The fitted model had an intercept of 4.66 and a coefficient of 1.39. The decision threshold estimated for expression scores was −1.56 Δlog(MFI) with variants having expression values less than this boundary classified as non-viable, and variants with expression scores greater than the boundary classified as viable. Greaney *et al*(*26*) used an expression threshold of −1 Δlog(MFI) to classify variants that have as large an expression deficit as mutations to core disulfide residues as nonviable. More recently, Greaney *et al*(*11*) used an expression threshold of −1.5 Δlog(MFI). Our estimated decision threshold is in-line with previous biologically-informed cutoffs.

We repeated the logistic regression analysis described above on the full Spike dataset from Dadonaite *et al*(*12*). In this dataset, only mutations seen in the pandemic were introduced to the full Spike in higher-order combinations. The dataset included 191,418 unique variants containing up to 38 mutations. For each variant, a functional score was estimated based on its relative frequency in the Spike-versus VSV-G-pseudotyped libraries. The fitted model had an intercept of 6.11 and a coefficient of 3.91. The decision threshold was estimated to be a functional score of −3.34, with variants having values lower than this cutoff classified as non-viable and variants with higher values classified as viable.

### Testing Neutralization

#### Sera Collection and Pooling

We collected 115 serum samples from individuals who were either convalescent, vaccinated, boosted, or experienced breakthrough infection (Supp Table S7). Samples were acquired from the POSITIVES study whose collections and primary outcomes have been reported elsewhere or through local Boston collaborations following institutional IRB approved protocols and with patient informed consent(*50*, *51*). For patients experiencing breakthrough infection, whole genome sequencing was performed on a nasal swab collected at the time of diagnosis to confirm infecting SARS-CoV-2 variant. Serum samples were acquired approximately 2-4 weeks after vaccination or convalescence. In order to maximize the number of designed Spike variant pseudoviruses that can be tested against the same immune serum panels, a total of 23 serum pools were created by grouping 5 individual patient samples per pool. The pooled sera are representative of the different populations that have existed throughout the pandemic: natural infection, primary vaccination, boosted, bivalent boosted, and breakthrough infection. When sufficient samples were available, individual serum samples were pooled based on neutralization titers resulting in low, medium, and high neutralization pools.

#### Neutralization Assay

Neutralizing activity against SARS-CoV-2 pseudovirus was measured using a single-round infection assay in 293T/ACE2 target cells. Pseudotyped virus particles were produced in 293T/17 cells (ATCC) by co-transfection of plasmids encoding codon-optimized SARS-CoV-2 Spike (containing G at position 614), packaging plasmid pCMV R8.2, and luciferase reporter plasmid pHR’ CMV-Luc. Packaging and luciferase plasmids were kindly provided by Dr. Barney Graham (NIH, Vaccine Research Center). The 293T cell line stably overexpressing the human ACE2 cell surface receptor protein was kindly provided by Drs. Michael Farzan and Huihui Ma (The Scripps Research Institute). For neutralization assays, serial dilutions of patient serum samples were performed in duplicate followed by addition of pseudovirus. Pooled serum samples from convalescent COVID-19 patients or pre-pandemic normal healthy serum (NHS) were used as positive and negative controls, respectively. Plates were incubated for 1 hour at 37 C followed by addition of 293/ACE2 target cells (1×10^4^ /well). Wells containing cells + pseudovirus (without sample) or cells alone acted as positive and negative infection controls, respectively. Assays were harvested on day 3 using Promega BrightGlo luciferase reagent and luminescence detected with a Promega GloMax luminometer. Titers are reported as the dilution of serum that inhibited 50% or 80% virus infection (ID_50_ and ID_80_ titers, respectively). We use the Wilcoxon signed rank test for comparing differences in neutralization.

#### Antigenic cartography

We use Racmacs(*52*) to perform multidimensional scaling (MDS) on our titration data measuring the neutralization sensitivity of all VOCs and designs for each serum pool. MDS is used to create a 2-dimensional representation of the pairwise antigenic distance between each VOC or design, as dictated by how each is neutralized by the different sera pools. MDS pairwise distances are consistent, which can be seen across pairs of variant-variant, variant-design, and design-design (Supp Fig S6). This allows us to visualize the trajectory of evolution in antigenic space, and showcase antigenic similarity between designs and later pandemic strains.

#### Nonhuman primate immunization and sera collection

The mosaic-8b and homotypic nanoparticles were described previously by Cohen *et al*(*7, 8*). Fourteen NHPs were primed with a mixture of nucleic acid-based SARS-CoV-2 vaccines 64-30 days prior to boosting with either a bivalent mRNA vaccine (n = 4), a mosaic-8b nanoparticle (n = 5), or a homotypic nanoparticle (n = 5) vaccine. Samples were collected 2-weeks post booster. The details of the immunization histories are described by Cohen *et al*.(*36*) The mixed immune history in these groups of pre-vaccinated NHPs is representative of a complex immune history in people who have been vaccinated and/or infected multiple times.

## Acknowledgements

The authors would like to thank Stephen Walsh, and Lindsey Baden at Brigham and Women’s Hospital (Boston, MA) for providing serum from vaccinated individuals for this study, and members of the Luban, Lemieux, Seaman and Marks labs.

## Data and Materials Availability

All code used for analysis in this study is publicly available on GitHub (https://github.com/debbiemarkslab/Vax_design). All data are available in the main text or the supplementary materials.

## List of Supplementary Materials

**Figure S1:**
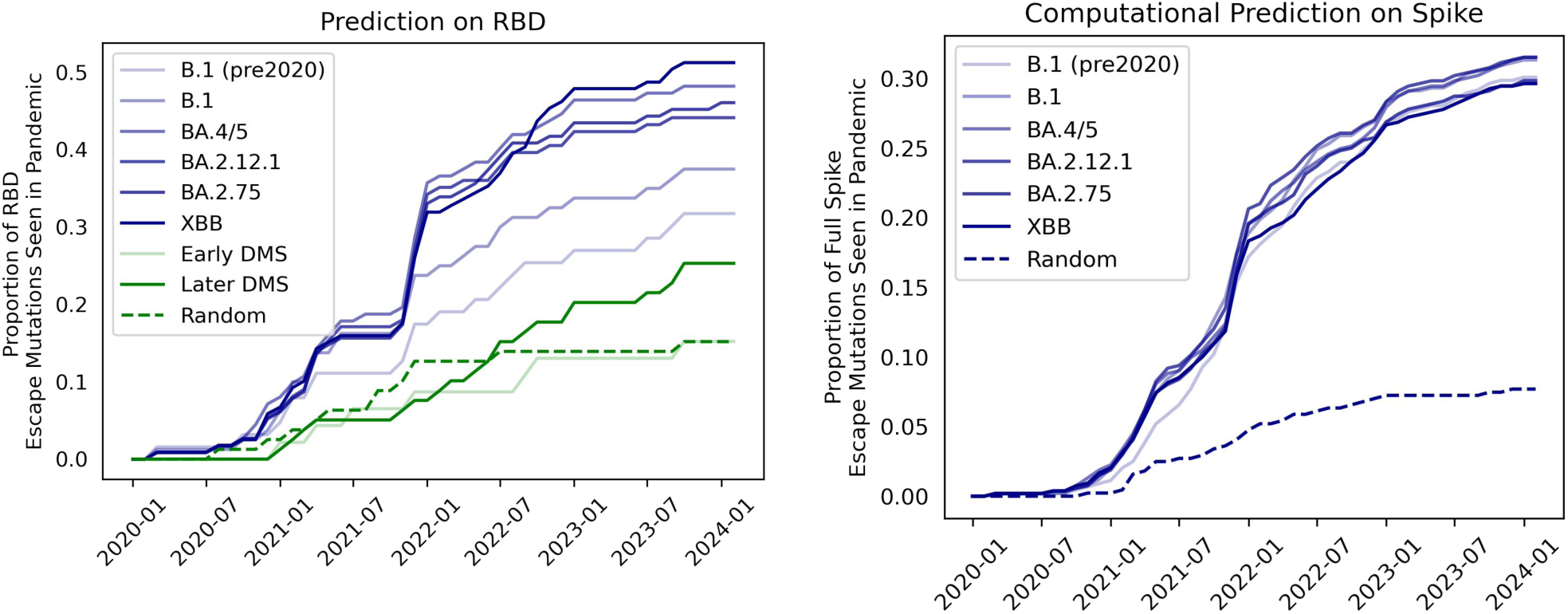
EVEscape better predicts later pandemic mutations when including earlier pandemic sequences in training. Proportions of escape mutations that have been seen more than 1000 times in GISAID as predicted by EVEscape (with and without SARS-CoV-2 sequences), mutational scans with pandemic antibodies, or randomly selected mutations in the RBD (left) or full Spike (right) protein (top 5%). EVEscape performance improves as pandemic sequences are incorporated into model training (blue lines based on variant model was trained up to). Analysis focuses on nonsynonymous RBD mutations that are a single nucleotide distance away from the ancestral sequence.

**Figure S2:**
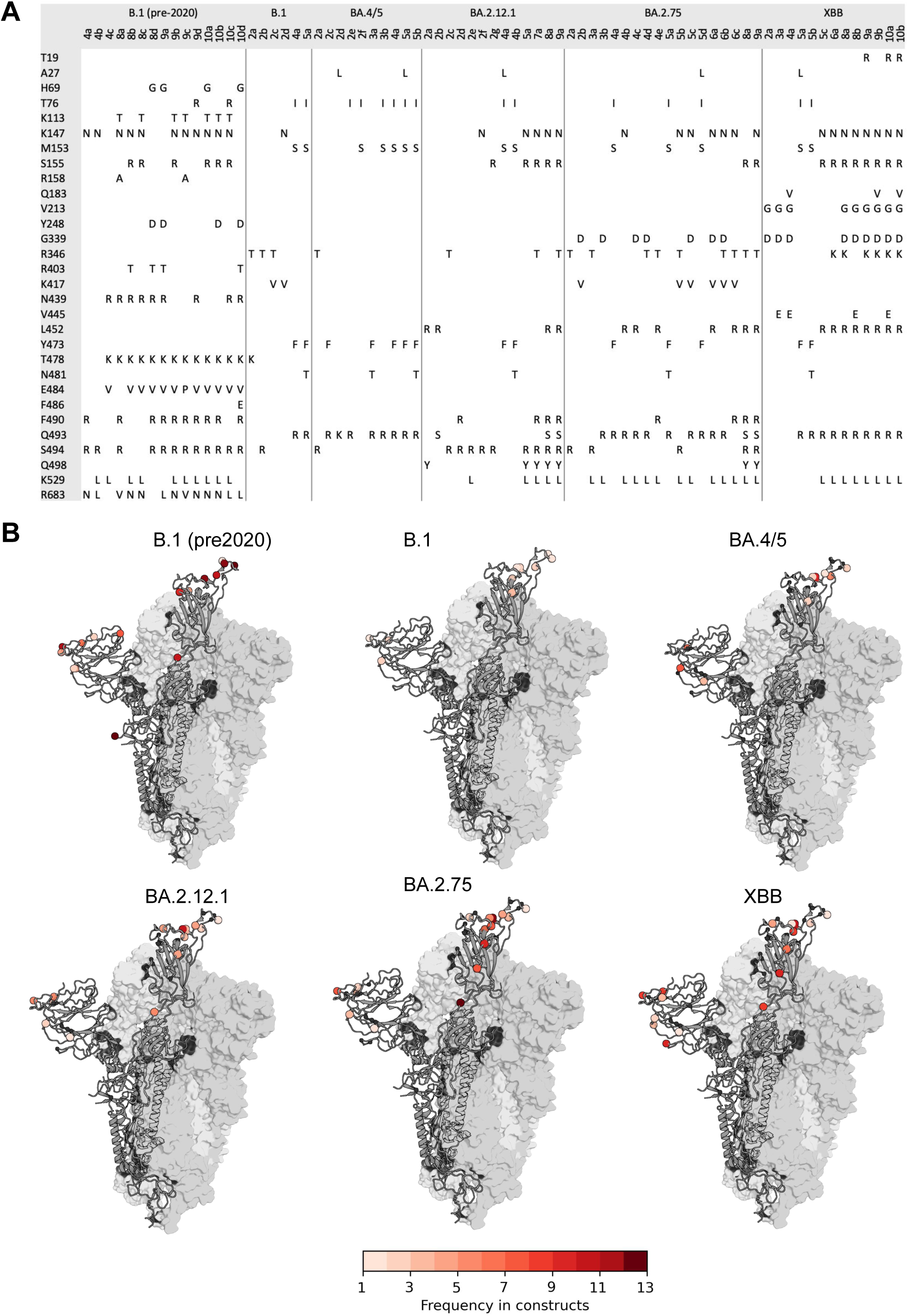
Overview of designed sequences. (A) Summary of designed spike sequences with two or more mutations on variant backgrounds (B.1, BA.4/5, BA.2.12.1, BA.2.75, and XBB). (**B**) Designed mutations across designed constructs in each of the five rounds mapped onto a representative Spike 3D structure (PDB ID: 7BNN), with coloring indicating the frequency with which a given residue was mutated in that round. Mutations in variant background sequences marked with small spheres.

**Figure S3:**
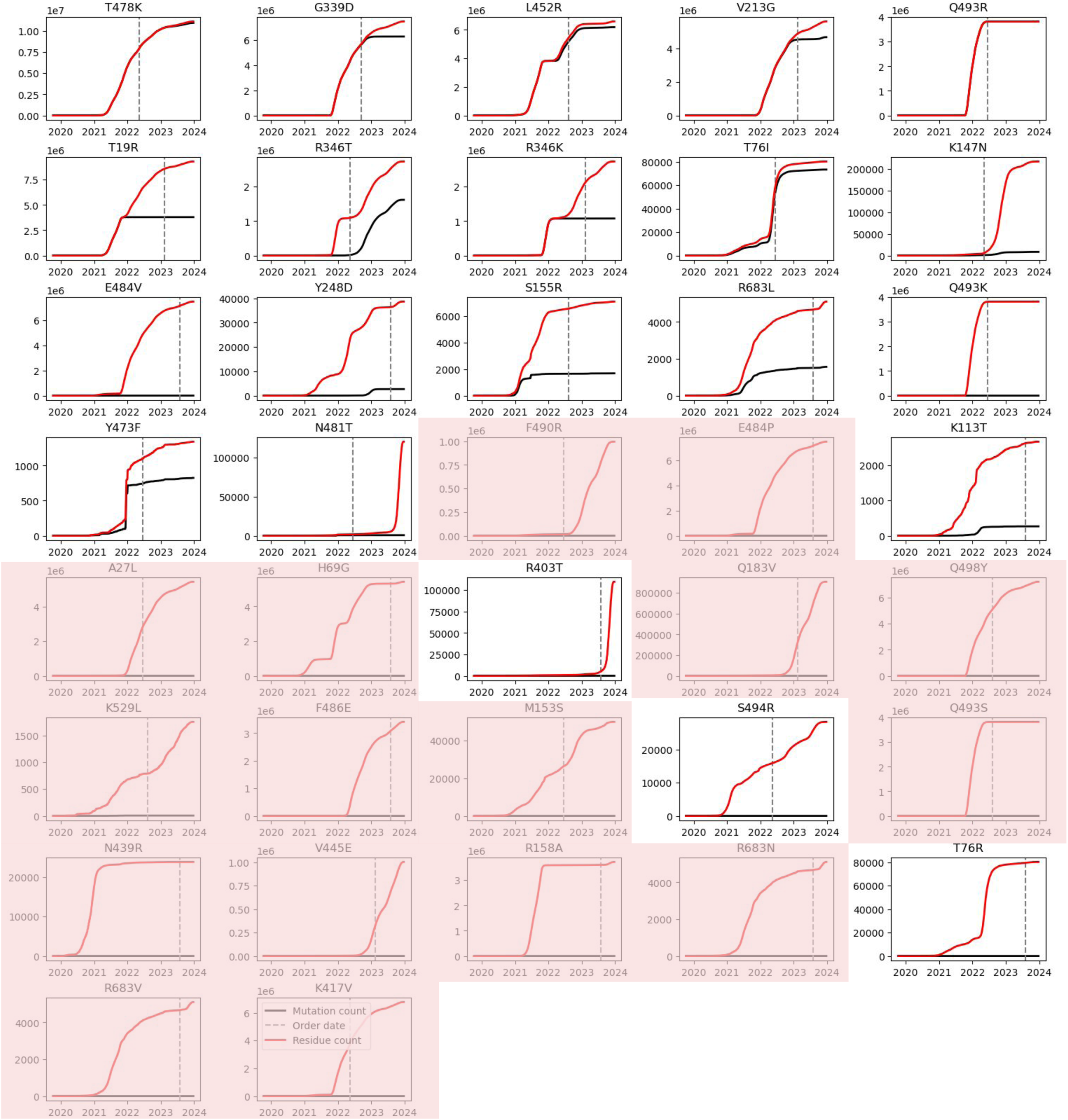
Count of designed mutation throughout the COVID19 pandemic. Black line shows cumulative mutation count in GISAID. Red line shows cumulative count of any mutations at that particular residue. Gray dashed line shows data construct containing that particular mutation was ordered. As expected, designed mutations which necessitate more than one nucleotide change (pink background), were rare in the pandemic. However, the corresponding residue positions were highly mutated during the pandemic, highlighting the model’s ability to identify mutable positions.

**Figure S4:**
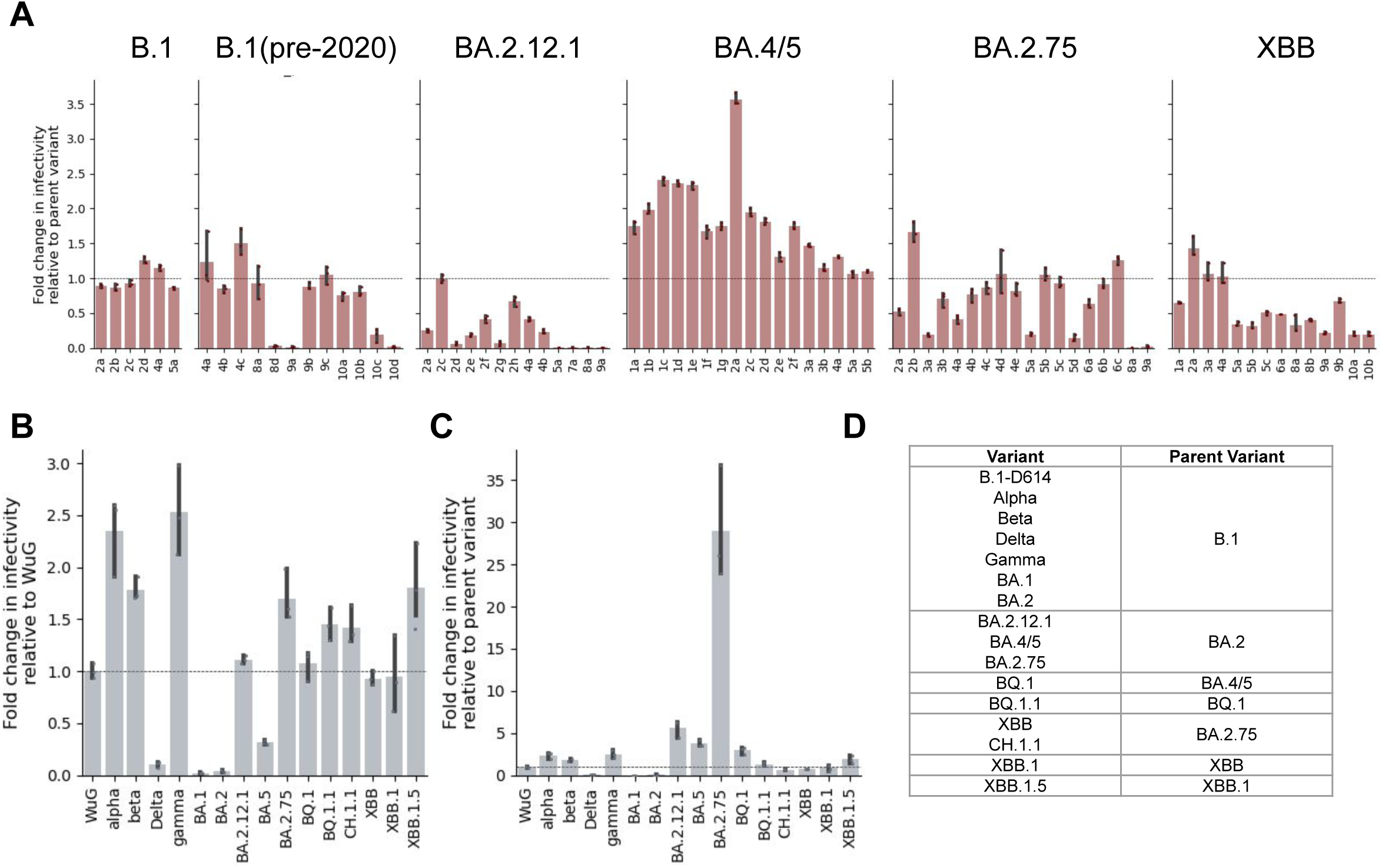
Infectivity Data. **(A)** Fold change in infectivity of designed constructs relative to the base variant on which they were designed. Error bars are 95% confidence intervals. **(B)** Fold change in infectivity of variants of concern (VOCs) relative to B.1. For consistency, we also compare the infectivity of the B.1. **(C)** Fold change in infectivity of variants of concern (VOCs) relative to their patient variant. **(D)** Summary of parent variants for VOCs.

**Figure S5:**
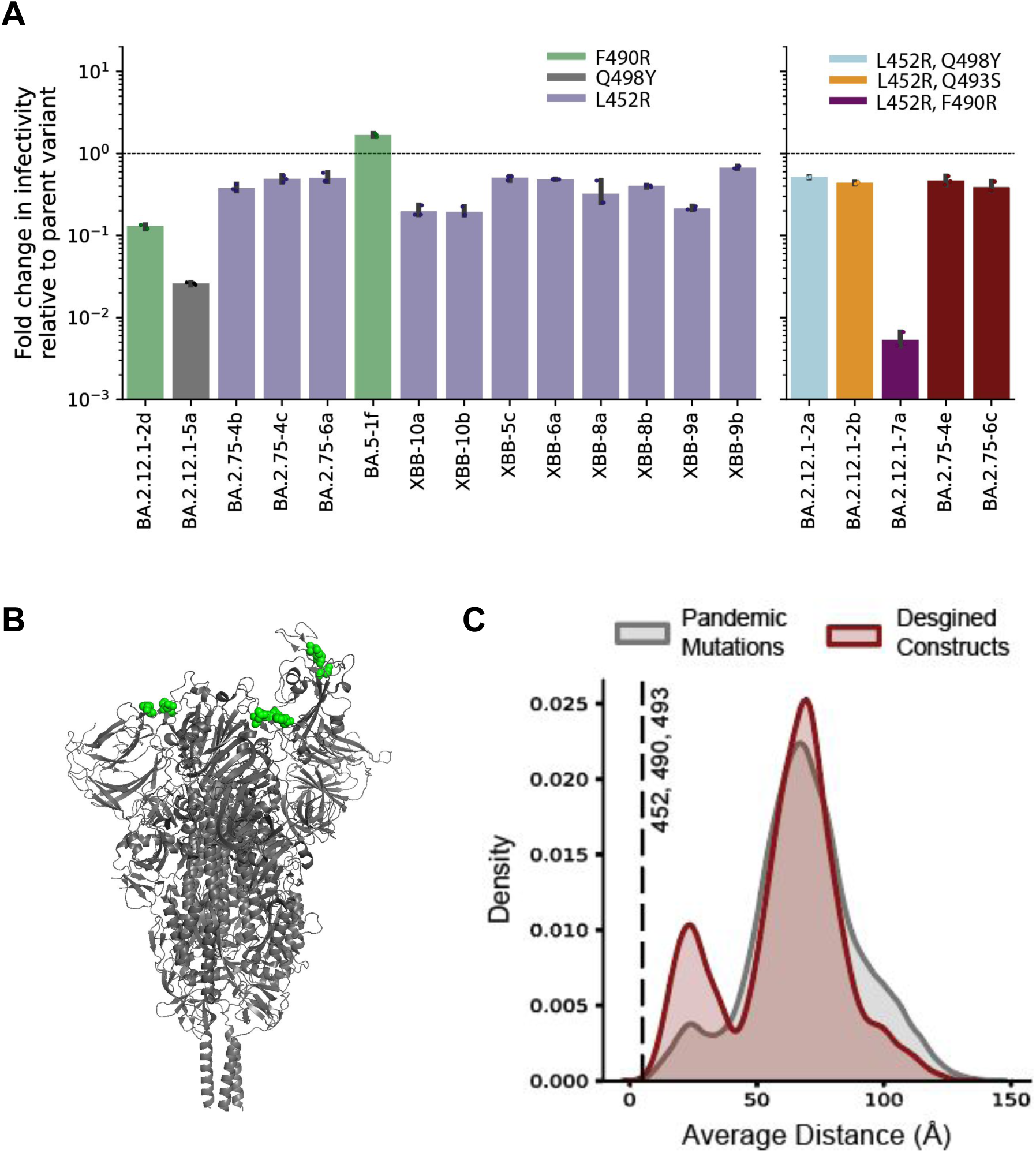
Proximity in three-dimensional structure suggests that the lack of infectivity may be explained by physical residue interactions. (**A**) Constructs containing some of these mutations individually, or in pairs, remain infectious. Error bars are 95% confidence intervals. (**B**) The four designed constructs which were not infectious shared a unique subset of mutations: L452R, F490R, Q493S, and Q498Y. Three proximal residues (452, 490, 493) are shown on Spike structure (PDB ID: 7BNN). (**C**) The three residues 452, 490, and 493 are closer in average distance than the average distance of any triplet of mutations in any pandemic strains or in any other designed constructs.

**Figure S6:**
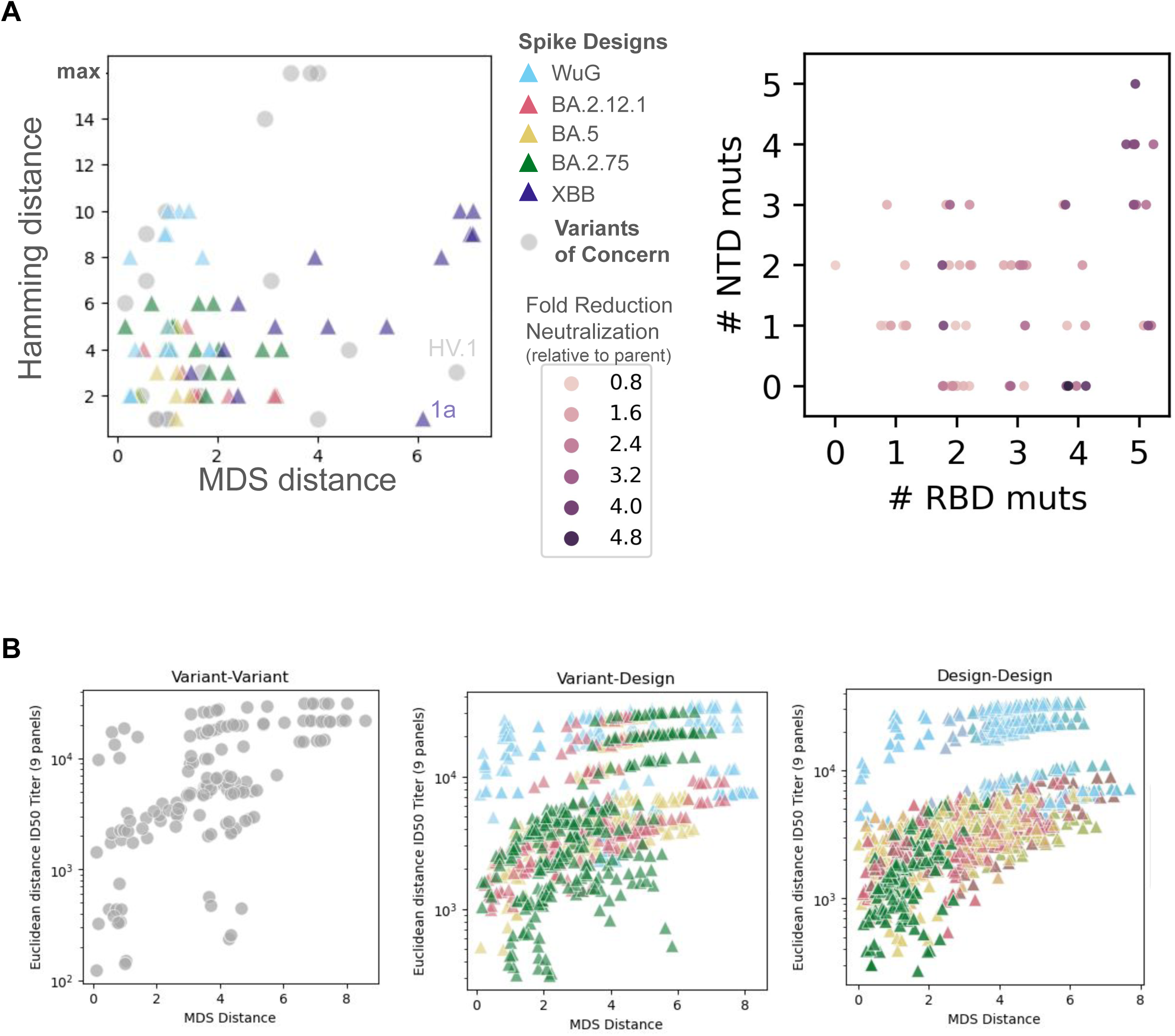
Antigenic distances are consistent and not entirely dependent on hamming distance. (**A**) Designs or pandemic variants and their distance by MDS (antigenic distance; Fig 3A) or hamming distance of Spike sequences from their parent variant (Fig 1C) (left). The maximum hamming distance shown is 16. Designs broken down by their number of NTD vs. RBD mutations, colored by their fold reduction in neutralization relative to their parent variant (right). Some designs or variants have outsized impacts on antigenic distance relative to their number of mutations (e.g., pandemic variant HV.1 and design XBB-1a). Designs are colored by their parent variant. (**B**) MDS distances are consistent with euclidean distances in ID50 titers across the 9 sera panels, when looking between all tested variant-variant, variant-design, and design-design pairs.

**Figure S7:**
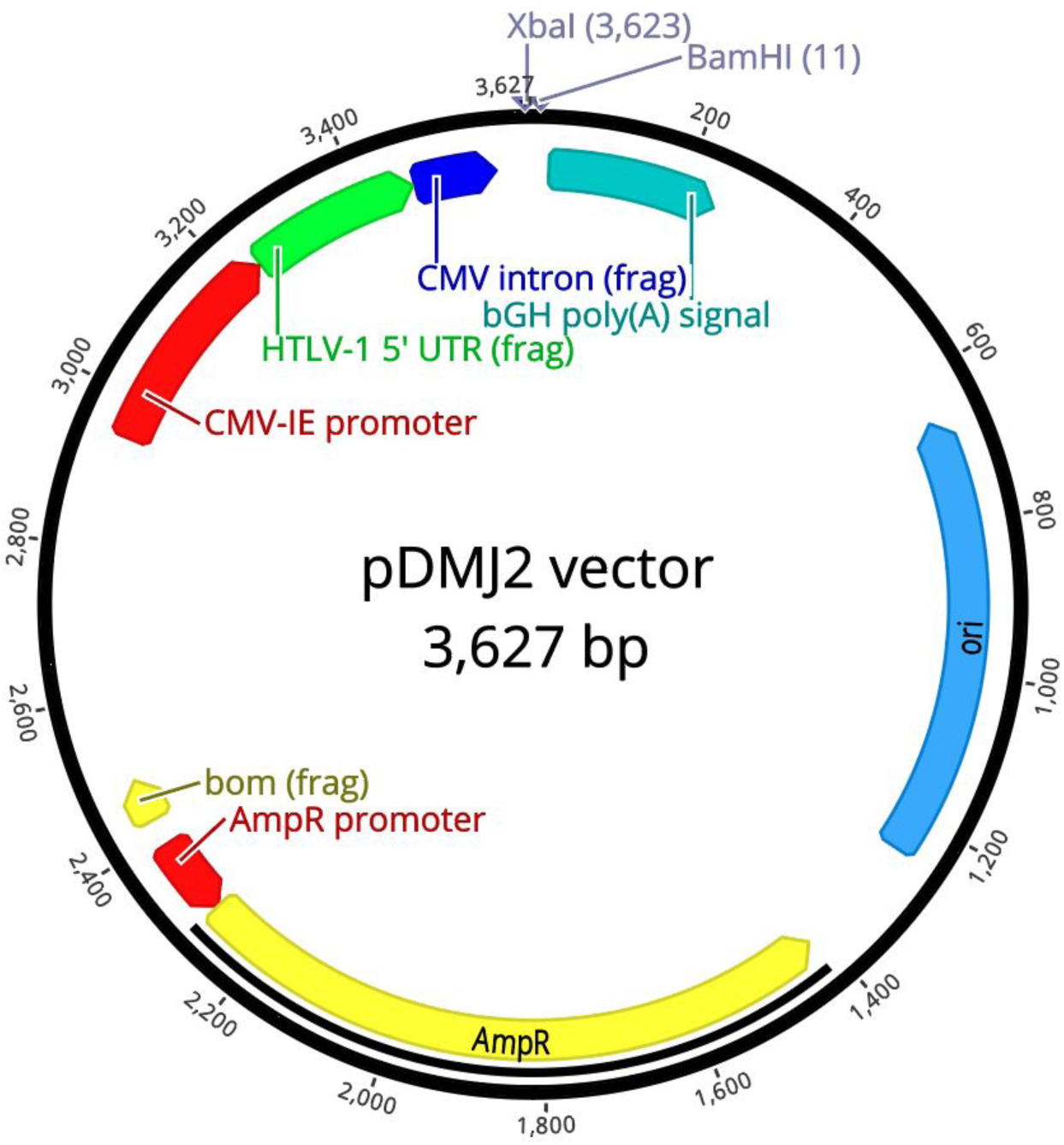
pDMJ2 plasmid map. (3,627 bp). Annotations: CMV-IE promoter: Cytomegalovirus immediate early promoter; HTLV-1 5’ UTR (frag): human T lymphotropic virus type 1 (HTLV-1) 5’ UTR fragment, CMV intron (frag): Cytomegalovirus intron fragment; bGH poly(A) signal: bovine growth hormone polyadenylation signal; ori: ColE1 origin of replication; AmpR: Ampicillin resistance gene; AmpR promoter: Ampicillin resistance promoter; bom (frag): ColE1 basis of mobility fragment. *XbaI* and *BamHI* restriction sites.

